# Identification of a persisting bacterial contamination of cell culture as *Brevibacterium*

**DOI:** 10.1101/2020.05.20.107698

**Authors:** P.P Mahesh, R. J Retnakumar, Sathish Mundayoor

## Abstract

Cell culture is an important prerequisite for many of the basic and applied researches in life sciences and contamination is a major problem faced by the researchers in maintaining healthy cell lines. We noted a dot like contamination in our THP1 cell line, which does not make the culture medium turbid as fast as other common contaminants and remains in the culture for many days before it makes the culture unusable. The contaminant was identified as a member of the genus *Brevibacterium* and we could find that the antibiotic rifampicin effectively suppresses the growth of the bacterium.

## Introduction

Human monocytic cell line THP1 was maintained in RPMI+10% FBS. A type of contamination previously thought as sediments from FBS or cell debri was detected, as it was observed increasing its number on repeated subculturing of the cell line and primarily assessed as a bacterium. The bacterium is non motile but it appeared vibrating, resembling to Brownian movement. A number of social forums discuss similar contamination detected from labs around the globe (https://www.researchgate.net/post/Dot-like_particles_in_cell_culture_is_it_contamination_or_not_No_change_in_pH_turbidity_cell_morphology_and_cell_growth,https://www.researchgate.net/post/Black_dots_like_contamination_in_my_cell_cultures_in_tightly_adhered_manner_What_to_do, http://www.protocol-online.org/biology-forums/posts/12608.html, https://www.youtube.com/watch?v=lo96fcXfcVs, https://www.youtube.com/watch?v=lQG_362L8VM).

The bacteria in low numbers do not affect the morphology of THP1 cells but prevent ideal differentiation of macrophages as observed under a light microscope. Also, there was no color change of the RPMI medium from brick red to yellow when the cells were maintained for many days. A study reports that the identification of *Achromobacter* in cell culture referring to a similar contamination but differs from our observation as *Achromobacter* is motile^1^. Another study reports dot like contaminating bacteria residing inside the cells and the study did not confirm the taxon of the bacterium^2^.

## Results and Discussion

Even though the bacteria do not affect the morphology of the cells, the rate of division and clustering of THP1 are largely reduced in the presence of bacteria. Washing the cells at low rpm centrifugation reduced the bacterial load but did not solve the problem. When the cells were maintained for weeks the rate of multiplication of the bacteria increased and ended up as large black colored aggregates and chains, and it made the cells no longer usable (Fig. 1a). Experiments demanding high cell density were practically impossible when the bacteria reached the aggregate forming stage. Therefore we decided to identify the bacterium by 16S rRNA sequencing. Since the bacteria did not grow on LB plates we attempted to make agar plates of RPMI+10%FBS and streaked with the cell culture supernatant when the bacteria reached the aggregate forming stage. Bacterial colonies appeared within a day (Fig. 1b), and colonies were scraped out and DNA was isolated by Phenol-chloroform method.

**Figure 1.**
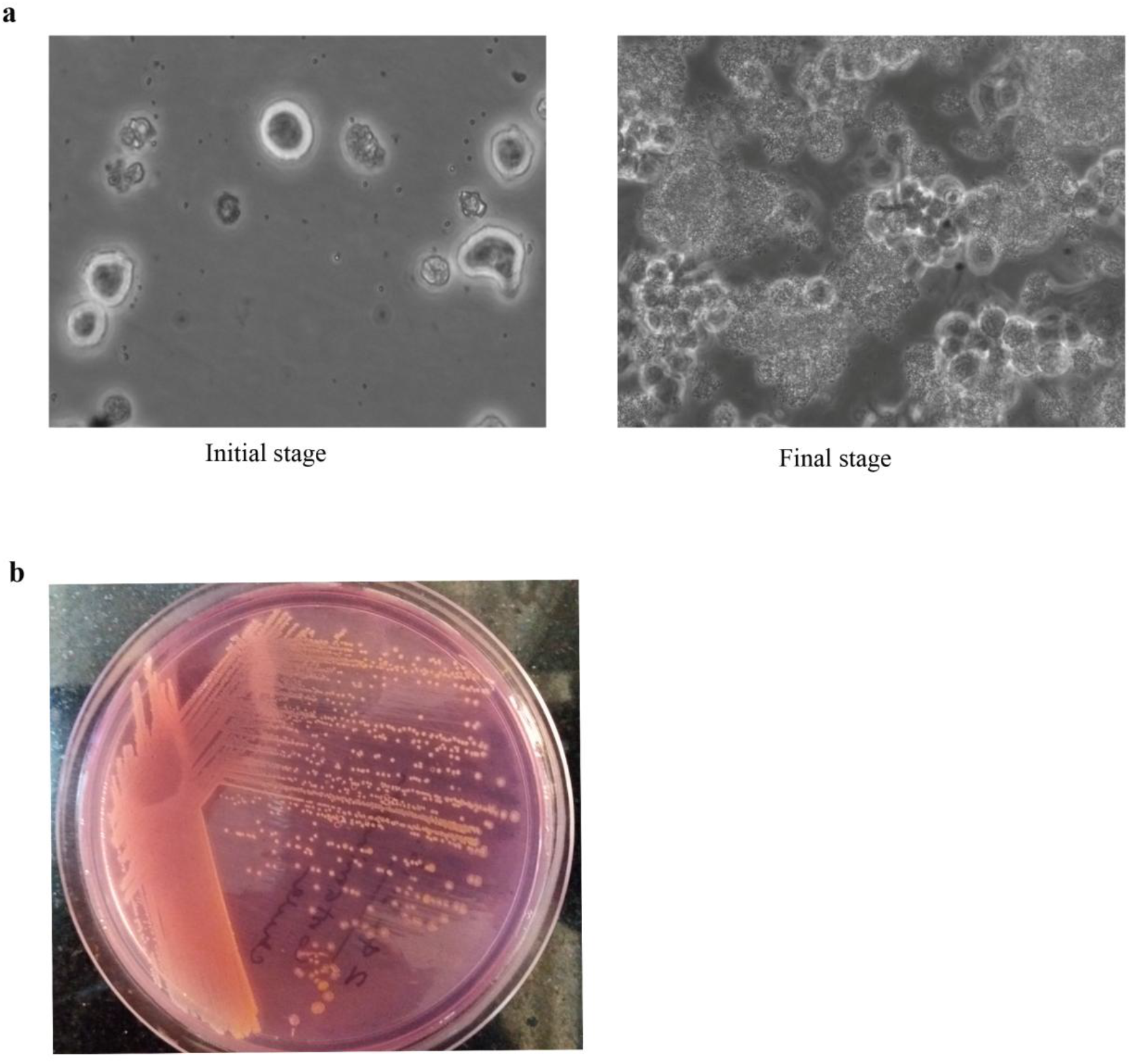
a) Initial and final stages of growth of the contaminant bacteria viewed under a light microscope (×200). b) Colonies of the bacteria grown on RPMI-FBS agar plate.

Initially we did 16S rRNA sequencing using a universal primer for bacteria which gave an amplification of <100bp stretch. When this sequence was submitted in NCBI Blast it returned different species but all of them were coming under Actinobacteria. Later we selected 2 sets of primers specifically designed for identification of Actinobacteria from a published study^3^. The primer pairs are listed in Table 1. A size of about 1300bp was amplified using each primer pair and processed for sequencing with respective primers. The maximum sequence length obtained was ~1099bp (Table 2).

**Table 1.**
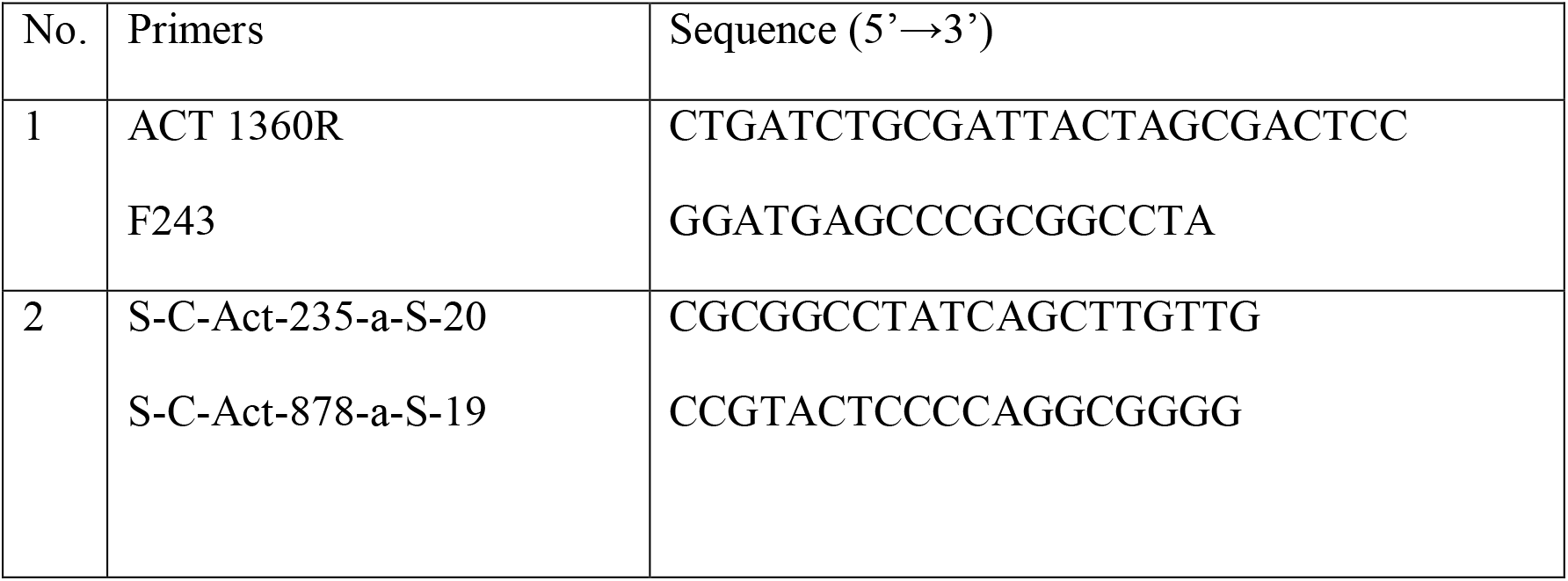
List of primers used for 16S rRNA sequencing

**Table 2.**
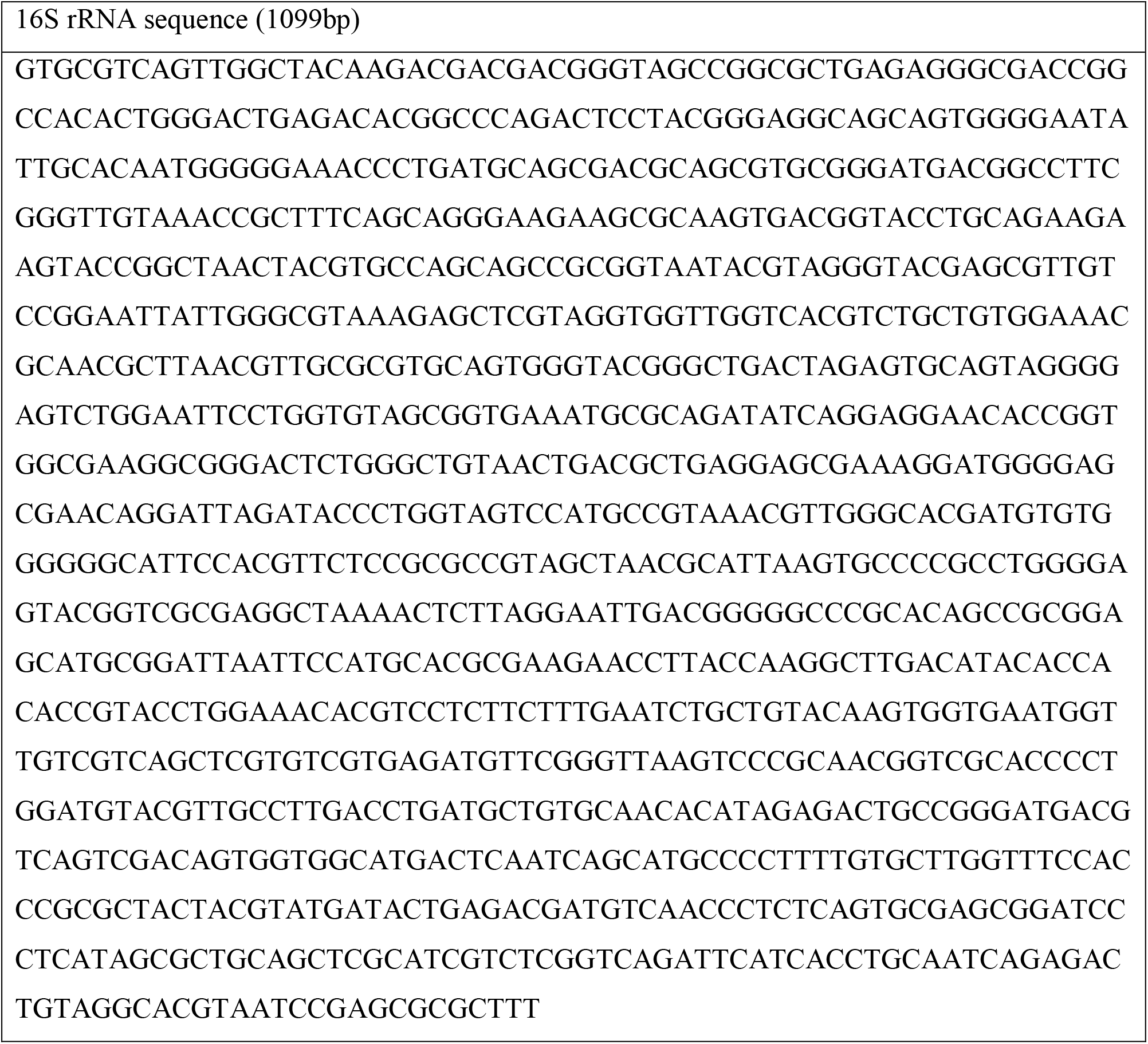
16S rRNA sequence

The sequence was submitted both in NCBI Blast and another database Ez Taxone to identify the bacterium. In both the cases the searches returned only one genus but different species of *Brevibacterium. Brevibacterium sanguini and Brevibacterium casei* were the top hits. Brevibacteria were reported as contaminants in milk products and found on human skin. If this is the case, it may be speculated that the source of contamination of cell culture was probably FBS or the contamination was from human handling.

Later we moved on to find out proper antibiotics to control the bacterium. A few antibiotics were selected for testing and most of them were actinomycete specific. The antibiotics selected were anti-bacterial anti-mycotic cocktail (Sigma) containing penicillin, streptomycin and amphotericin; ciprofloxacin, piperacillin, chloramphenicol, kanamycin, erythromycin, rifampicin, linezolid, vancomycin, amoxicillin, cotrimoxazole, meropenem and cefazoline. Antibiotics other than the cocktail (used at 1-4X), were tested at 1-10μg/ml.

Among the antibiotics tested rifampicin at 10μg/ml gave the most effective result. Cotrimoxazole and ciprofloxacin resulted in considerable reduction in the rate of division of THP1 cells at all concentrations tested. At the same time none of the antibiotics tested removed the bacteria completely. Treatment of any of the antibiotics at >10μg/ml did not improve the result significantly. Also, different combinations of antibiotics did not improve the results compared to what was obtained when they were used alone. Rifampicin at 10μg/ml increased the rate of division and clustering of THP1, resulted in quicker color change of the medium than the contaminated cultures and showed better differentiation of macrophages by visual analysis. Culturing of THP1 in the presence of rifampicin at 10μg/ml prevented the bacteria from entering to the aggregate forming stage (possibly log phase of the bacterium) and the long time culturing of cells (at least one month) with the antibiotic did not cause any considerable change in cell division or cell health as assessed by morphological analysis. When rifampicin was withdrawn, the bacteria did not enter into log phase for at least 3 days, possibly because the antibiotic might have extended the lag phase of the bacteria. Even though rifampicin used at 10μg/ml did not cause any considerable change in the morphology of THP1 cells, there are reports on the side effects of the antibiotic on mammalian cells when it was used at higher concentrations^4,5^.

## Materials and methods

### Cell culture

THP1 monocytic cell line was maintained in RPMI-1640 medium (R4130, Sigma) supplemented with 10 % FBS. For obtaining macrophage monolayer THP1 cells with required cell density were treated with 20ng/ml of PMA (P8139, Sigma) and after one day PMA was removed with 2 washes of RPMI.

### l6SrRNA sequencing

The bacterial colonies obtained on RPMI-FBS agar was used to isolate genomic DNA by standard phenol-chloroform method. 16SrRNA gene was amplified using two sets of primers – 16SrRNA universal primer and specific primer for Actinobacteria. Sequencing PCR was carried out by BigDye Terminator chemistry and the amplicons were sequenced using Sanger sequencing. Analysis of the resulting sequences by NCBI Blast analysis and Ez Taxone/ EzBioCloud database (ezbiocloud.net) revealed that the isolated bacteria belong to *Brevibacterium* species, with *Brevibacterium sanguini* and *Brevibacterium casei* being top hits.

### Antibiotics

Antibiotics used for the treatment include anti-bacterial anti-mycotic cocktail (A5955, Sigma), chloramphenicol (C0378, Sigma), kanamycin (K1377, Sigma) and rifampicin (R3501, Sigma). Other antibiotics used are pharmaceuticals.

## Acknowledgements

This work was supported by intramural fund from RGCB. PPM was supported by a fellowship from Council for Scientific and Industrial Research, Govt of India.

## Author Contribution statement

PPM, RJR and SM designed the study. PPM and RJR performed the research. PPM and SM wrote the manuscript together.

## Competing interests

The authors declare no competing interests.

## Notes

### Competing Interest Statement

The authors have declared no competing interest.

